# HSV-1 Oncolytic Virus Targeting CEACAM6-Expressing Tumors Using a Bispecific T-Cell Engager

**DOI:** 10.1101/2023.01.02.522257

**Authors:** Yanal M. Murad, I-Fang Lee, Xiaohu Liu, Zahid Delwar, Jun Ding, Guoyu Liu, Olga Tatsiy, Dmitry Chouljenko, Gregory Hussack, Henk Van Faassen, William Wei-Guo Jia

## Abstract

VG21306 is a novel oncolytic virus (OV) that encodes a secretable bispecific T-cell engager targeting Carcinoembryonic antigen cell adhesion molecule 6 (CEACAM6)-expressing tumors. Delivering a T-cell engager locally to a tumor mass will circumvent physical barriers that prevent antibodies from penetrating the tumor, and will mitigate the off-tumor, on-target toxicity risk. Both in vitro and in vivo testing demonstrated the expression of a functional T-cell engager capable of binding both targets. The efficacy of the engager was demonstrated in vitro, where addition of the engager payload to the OV enhanced anti-tumor efficacy against tumor cells overexpressing CEACAM6. Moreover, we have demonstrated the engager’s ability to induce bystander killing in cells lacking CEACAM6 expression, as well as engaging exhausted T cells and inducing tumor cell death. The safety of the engager was demonstrated by the lack of binding to normal human tissue or normal tissue adjacent to tumors, as well as the absence of any measurable leakage of the expressed engager into the blood of mice treated by intratumoral OV injection.

## Introduction

Redirecting the activity of T cells to target tumor cells is a powerful tool for cancer immunotherapy. The idea of an artificially constructed bispecific antibody was born almost simultaneously with the first insights into an antibody’s structure and function by Rodney Porter, who won the Nobel Prize for reporting on the basic structure of the immunoglobulin molecule [1]. The idea gained traction in the 1980s when a slew of reports and research work reported on using bispecific T-cell-engager (BiTE) antibodies to redirect the immune response against a specific target or antigen. This interest culminated in 2014 when the FDA granted breakthrough therapy status to blinatumomab for the treatment of acute lymphoblastic leukemia (ALL) [2].

Since the approval of the first BiTE, multiple tumor antigens were targeted by T-cell engagers, including epithelial cell adhesion molecule (EpCAM) [3][4][5], human epidermal growth factor receptor (HER) family [6], carcinoembryonic antigen (CEA) [7][8], and prostate-specific membrane antigen (PSMA) [9]. However, the use of the aforementioned BiTEs have been associated with severe, potentially fatal toxicities, while the efficacy against solid tumors has been limited, owing to physical barriers and an immunosuppressive tumor microenvironment [10]. Thus, there is a need for an improved therapeutic strategy that ameliorates BiTE-mediated toxicity during cancer therapy.

Oncolytic viruses (OVs) have been engineered to express and deliver multiple payloads to tumors [11]. In particular, HSV-1-based OVs are known to have high capacity to accommodate multiple payloads [12][13]. The virus may be engineered to express a variety of immunomodulators—such as IL-12, IL-15, and IL-15RA1—and it may be modified to enable tumor-specific virus replication and/or tumor-specific payload expression. Multiple reports have described OVs that carry BiTEs as payloads [14][15].

VG21306 is the first OV that carries a T-cell engager payload targeting CEACAM6 antigen. CEACAM6 is overexpressed in multiple tumors, including in pancreatic [16], [17], lung [18], and other types of cancer [19]. Several antibodies have been developed to target this antigen [20]–[23][24][25]. In this paper, we report the use of a novel anti-CEACAM6 single-domain antibody to generate a T-cell engager (also known as UCHT-2-03) targeting CEACAM6 antigen [26], along with UCHT1, an anti-CD3 antibody known to target T cells [27]. The use of an OV to express a bispecific T-cell engager combines two complementary cancer immunotherapy approaches to synergistically enhance antitumor activity. While an OV can infect and lyse a small number of cells in the tumor mass, it serves to prime the tumor microenvironment for immunotherapy by increasing the levels of tumor-infiltrating lymphocytes. Moreover, the lytic destruction of a subset of tumor cells will cause the release of tumor-associated antigens (TAAs) that may facilitate an anti-tumor adaptive immune response. Viral infection in the tumor microenvironment may also provide a danger signal that partially counteracts the immunosuppressive tumor microenvironment. On the other hand, the BiTE will take advantage of the increased infiltration of T cells into the tumor induced by the OV treatment, which we have demonstrated in previous work [28], [29], promoting T-cell activation and tumor-cell killing in the presence of cells displaying a selected TAA on the cell surface.

Consequently, the OV expressing a tumor-specific T-cell engager would be preferentially used to treat tumors that overexpress the selected TAA used for the tumor-targeting function of the T-cell engager.

This report summarizes the results of in vitro and in vivo testing of this novel OV, and provides a mechanistic study to elucidate the enhanced efficacy over the backbone virus lacking the engager payload.

## Materials and Methods

### Cell lines

African green monkey kidney (Vero) cells, the human tumor cell lines LS174T, MDA-MB-468, MDA-MB-231, BxPC3, and HepG2, were obtained from the American Type Culture Collection (Manassas, VA, USA). Cells were maintained in Dulbecco’s Modified Eagle’s Medium (DMEM) (Gibco-BRL, Grand Island, NY, USA) supplemented with 10% fetal bovine serum (FBS) (Thermo Fisher Scientific, Waltham, MA, USA). A549 cell line stably expressing CRISPR Cas9 nuclease (Genecopoeia) was used to generate CEACAM6 knock-out cells (A549-CEACAM6-KO) using lentivirus particles carrying the CRISPR human sgRNA for CEACAM6 (Genecopoeia). Clones were selected using puromycin and the CEACAM6-expression knockout was confirmed by flow cytometry and Western blot.

### Construction, expression and characterization of UCHT-2-03 engager

The T-cell engager (UCHT-2-03) was cloned and expressed using ExpiCHO cells (Invitrogen), and the recombinant protein was purified using Ni-column and size-exclusion chromatography (SEC).

The purity of the proteins and their aggregate formation or lack thereof were assessed using a Superdex™ S200 10/300GL SEC column (Cytiva, Vancouver, Canada) and an ÄKTA FPLC™ system (GE Healthcare) in HBS-EP buffer (10 mM HEPES, 150 mM NaCl, 3 mM EDTA, 0.005 % Tween 20, pH 7.4) at a flow rate of 0.8 mL/min. The binding affinity of the UCHT-2-03 engager to human CEACAM6 (Acro Biosystems, Newark, DE; Cat# CE6-H5223) and human CD3δ/ε heterodimer (Acro Biosystems; Cat# CDD-H52W1) was determined through surface plasmon resonance (SPR) using a Biacore T200 instrument (Cytiva) at 25 °C in HBS-EP running buffer. CEACAM6 and CD3δ/ε heterodimer, as well as human EGFR (GenScript, Piscataway, NJ; Cat# Z03194) as a control surface, were immobilized onto a CM5 Series S sensor chip (Cytiva) using standard amine coupling in 10 mM acetate buffer, pH 4 (Cytiva). A surface blocked with ethanolamine served as the reference flow cell. The SEC-purified UCHT-2-03 engager was injected over all surfaces at a flow rate of 40 μL/min, with 180 s of contact time and 300 s (CEACAM6, EGFR) or 1800 s (CD3) of dissociation time. The concentration range of UCHT-2-03 injected was 0.25 – 4 nM (CD3) and 0.625 – 10 nM (CEACAM6, EGFR). Regeneration was performed with 10 mM glycine, pH 1.5, for 30 s at 50 μL/min (CD3) or HBS-EP running buffer (CEACAM6). Reference flow cell subtracted sensorgrams were fit to a 1:1 binding model using BIAevaluation software v3.2 (Cytiva) to calculate kinetics and affinities (n = 3).

A second SPR assay was performed to demonstrate simultaneous co-engagement of UCHT-2-03 with CEACAM6 and CD3, using HBS-EP running buffer and a flow rate of 40 μL/min. UCHT-2-03 (10 nM) was injected over CEACAM6 and CD3 surfaces for 150 s. This was followed immediately by a second injection of CD3 (10 nM), CEACAM6 (10 nM) or HBS-EP buffer for an additional 150 s. Surfaces were regenerated as described above.

### Enzyme-Linked Immunosorbent Assay (ELISA)

A sandwich ELISA was developed to measure the concentration of UCHT-2-03 engager in culture medium as well as in the serum of mice treated with VG21306 to measure the leakage of the engager into the blood. A 96-well Immuno Maxisorp flat-bottom plate (Thermo Fisher Scientific) was coated overnight with 100 ng/well (in 100 μL/well volume) of purified human CEACAM6 protein (Sino Biological). Standards containing known concentrations of the UCHT-2-03 recombinant protein or test samples were applied to the plate for 1 hour, followed by applying 100 ng/well (in 100 μL/well volume) of Flag-tagged human CD3δ/ε heterodimer (Sino Biological) in 100 μL of Dulbecco’s phosphate-buffered saline (DPBS). After a 1-hour incubation, horseradish peroxidase (HRP)-conjugated anti-Flag antibody was applied to the plate. The binding of target was detected via 3,3’,5,5’-tetramethylbenzidine (TMB) substrate. Absorbance measurements were collected at a wavelength of 450 nm on a microplate reader (Molecular Devices, San Jose, CA, USA).

### Recombinant virus construction

All viruses used in this study were constructed using herpes simplex virus type 1 (HSV-1) strain 17 as the backbone, and viral mutagenesis was performed using standard lambda Red-mediated bacterial artificial chromosome (BAC) recombineering techniques in *Escherichia coli.* The construction and characterization of recombinant HSV-1 OVs, including a transcriptional and translational dual-regulated (TTDR) virus similar to VG21306 was published previously [30][12]. Briefly, the native ICP27 promoter was replaced with the tumor-specific CEA promoter and the terminal repeat region was deleted to remove one of the two copies of ICP34.5, ICP0, and ICP4. Five tandem repeats of binding sites with perfect complementarity to microRNA (miR)-124 and miR-143 were inserted into the 3’-untranslated region of the remaining single copy of ICP34.5. The 28 C-terminal amino acids of glycoprotein B were also deleted, and an expression cassette for human IL-12, IL-15, and IL-15 receptor alpha (IL-15Ra) subunit isoform 1 driven by the tumor-specific CXCR4 promoter—and with each element separated by selfcleaving P2A peptides—was inserted between viral genes UL3 and UL4. As for the T-cell engager sequence, it was cloned between the viral genes US1 and US2 and was driven by the EF1α promoter.

Additional recombinant HSV-1 controls were constructed for testing purposes, including VG1905—which lacks the cytokines and T-cell engager payload of VG21306, but is otherwise identical—and VG2003, which only lacks the T-cell engager, but retains all other elements from VG21306.

The QIAGEN HiSpeed Plasmid Midi Kit was used to isolate mutant BACs, and virus was recovered in Vero cells following transfection using Lipofectamine™ 2000. Genomic integrity of recombinant HSV-1 was verified by restriction enzyme digestion and by targeted sequencing of all modified genomic regions.

### UCHT-2-03 engager cross-reactivity characterization

The engager cross-reactivity with normal human tissue, as well as with tumor tissues, were evaluated in multiorgan tumor tissue microarray slides containing 54 cases of tumor of varieties of organ tissue, plus 18 normal tissue or adjacent normal tissue, and a multiple organ normal tissue microarray containing 24 cases of normal tissue, divided into two identical 48-core arrays (US Biomax, Inc.) using deparaffinized sections. Antigen retrieval was performed using 2100 Retrieval kit, and His-tagged UCHT-2-03 was used to stain the slides, with biotinylated anti-6X His-tag antibody (abcam) used as a secondary detection antibody. VECTASTAIN^®^ Elite^®^ ABC-HRP Kit, Peroxidase (Vector Laboratories) and 3,3’-diaminobenzidine tetrahydrochloride hydrate (DAB) (Sigma) were used for the final development of color. For scoring the intensity of staining, ordinal scores (e.g., 0, 1, 2, 3, and 4) are reflective of cellular immunostaining frequency or intensity. Using frequency, the immunostaining incidence (%) was estimated in each tissue, which defined the respective score (e.g. “0”: none, “1”: 1 - 25%, “2”: 26 - 50%, “3”: 51 - 75%, and “4”: 76 - 100% stained). For intensity, the immunostaining score was evaluated (e.g., “0”: none, “1”: weak, “2”: moderate, “3”: strong, and “4”: very strong).

### In vitro engagement of tumor cells and T cells by UCHT-2-03 engager

CEACAM6-positive NSCLC A549 tumor cells as well as A549-CEACAM6-KO cells were stained with CellTracker™ Red, seeded into a 24-well plate, and incubated at 37°C for 24 hours. Next, pan-T cells were stained with CellTracker™ GFP and then co-cultured with previously seeded A549 or A549-CEACAM6-KO cells. Cells were then incubated at 37°C with either DPBS or VG21306 or VG2003 virus (virus expressing IL-12 and IL-15/IL-15RA1) at an MOI of 2 for 18 - 20 hours. Cells were then washed twice with DPBS and fixed with 4% paraformaldehyde (PFA). Fixed cells were then stained with anti-HSV-1 antibody to detect virus-infected cells. Imaging (20X) was performed using a fluorescence microscope.

### In vitro killing effect of UCHT-2-03 engager

CEACAM6-positive A549 and MDA-MB-468 tumor cells were labelled with 0.3 mM of CellTrace™ Far Red (Invitrogen), seeded into a 96-well, flat-bottom plate (2 x 10^4^ cells/well), and co-cultured with human pan-T cells from a healthy donor (STEMCELL Technologies) at an Effector:Target ratio of 10:1 in the presence of purified UCHT-2-03 T-cell engager at 37°C for 48 hours. Cytotoxicity of CellTrace™ Far Red-labelled tumor cells was measured by 7-AAD staining (Invitrogen) and flow cytometry. The percentage of tumor cell cytotoxicity was calculated as [% 7-AAD^+^ cells ÷ (% 7-AAD^+^ cells + % 7-AAD^-^ cells)] x 100.

### Engager-enhanced cytotoxicity

A549 tumor cells (1.5 x 10^4^ cells/well) were seeded into a 96-well, real-time cell analysis (RTCA) E-plate and incubated at 5% CO_2_ and 37°C overnight. Seeded tumor cells were subsequently co-incubated with MOI=0.1 of VG21306 (virus expressing IL-12, IL-15/IL-15RA1, and UCHT-2-03 T-cell engager), VG2003 (virus expressing IL-12 and IL-15/IL-15RA1), or VG1905 (backbone virus without payloads) virus and human peripheral blood mononuclear cells (PBMCs) isolated from a healthy donor. A549 cells with human PBMCs in the presence of purified UCHT-2-03 engager protein was also included as a control. The cytolysis of A549 cells was monitored over a period of 5 days using xCELLigence^®^ RTCA eSight device (Agilent Technologies, Inc., Santa Clara, CA) by impedance.

### In vivo T cell activation by UCHT-2-03 engager

Experimental animal procedures were approved by the BRI Biopharmaceutical Research Inc. Animal Care Committee (Vancouver, BC, Canada) and followed the guidelines and policies of the Canadian Council on Animal Care (CCAC).

A549 tumor cells (5 x 10^6^ A549 cells) and human PBMCs from a healthy donor (5 x 10^6^ A549 cells) were inoculated into NOD scid gamma (NSG) mice on Day 0. At Day 9, all mice were randomly divided into three groups and intratumorally injected with vehicle alone, 5 x 10^7^ PFU of comparator VG2003 virus, or 5 x 10^7^ PFU of VG21306 virus. Tumor samples were harvested at 72 hours post injection and stained with anti-human CD45, CD3, CD4, CD8, and CD69 antibodies. The activated total CD3^+^, CD8^+^, and CD4^+^ T cells in tumors were detected and analyzed by flow cytometry.

### Testing bystander killing of UCHT-2-03 engager

To test the bystander killing effect of UCHT-2-03 engager, BxPC3 cells (CEACAM6-positive cells) and HepG2 (CEACAM6-negative cells) were labelled with either carboxyfluorescein succinimidyl ester (CFSE) (BxPC3) or CellTrace™ Far Red (HepG2), then incubated at 37°C for 48 hours either separately or in co-culture (1:1 ratio) with the UCHT-2-03 engager (concentrations ranging from 0 - 10 ng/mL) and pan-T cells. Cell killing was detected using flow cytometry by 7AAD assay. Also, granzyme B and IFN-γ secretion were measured using ELISA as described by the manufacturer (R&D Systems).

### T-cell exhaustion assay

Pan-T cells (STEMCELL Technologies) were subjected to continuous stimulation as described by Dunsford et. al [31] with modifications. Briefly, T cells were cultured in ImmunoCult™-XF T Cell Expansion Medium (STEMCELL Technologies) at 2.5 x 10^6^ cells/mL and ImmunoCult™ Human CD3/CD28/CD2 T Cell Activator (STEMCELL Technologies) at 25μL/mL. The medium was changed every 2 - 3 days, with fresh CD3/CD28/CD2 T Cell Activator added at the recommended dose. The medium was changed three times for a total of four stimulations (9 days). On Day 9, stimulated cells were collected and resuspended in RPMI 1640 medium with 10% FBS prior to assay setup.

Co-culture assays: Unstimulated T cells were thawed and were allowed to rest overnight post thaw. A549 cells were seeded overnight at 1.2 x 10^4^ cells/well, and 0.5 nM or 0.05 nM of engager proteins and stimulated or unstimulated T cells at 10:1 or 5:1 Effector to Target ratio were added to the culture. The plates were then incubated at 37°C for 48 hours then analyzed by flow cytometry and ELISA or monitored for target cell death using xCelligence eSight (Agilent Technologies, Inc., Santa Clara, CA).

The co-culture assays for stimulated T cells were carried out after 9 days of stimulation in a similar way as described above.

Flow cytometry: The stimulation of T cells was monitored on Day 1 and Day 9 of stimulation with Attune NXT Flow Cytometer. The T cells were analyzed for the expression of CD3 (BioLegend), CD4 (APC-eFluor780, Invitrogen), CD69 (BV786, BD Biosciences), NKG2D (PE, Invitrogen), Tim-3 (BV510, BioLegend), LAG-3 (BV605), TIGIT (AF700, Invitrogen), and PD-1 (APC, BioLegend). The cells were also stained for viability using 7AAD (Invitrogen). The co-cultured T cells were also analyzed by flow cytometry after 48 hours with the antibodies indicated above, and the supernatant was used in a human IL-2 ELISA (Invitrogen), human IFN-gamma ELISA (Invitrogen) and granzyme B ELISA (R&D Systems) after 48 hours of incubation at 37°C, following kit instructions.

PD-L1 expression on tumor cells: A549 cells incubated with stimulated T cells were also analyzed for PD-L1 expression (APC, Invitrogen) and viability with 7AAD staining.

### Leaking of engager protein in A549 and HepG2 tumor models

CEACAM6-positive A549 tumor cells (2.5 x 10^6^ cells in 200 μL) or CEACAM6 knockout A549 cells were inoculated subcutaneously into the right flank of athymic nude mice. Once tumors reached ~100 mm^3^ in size, mice were randomly assigned to respective treatment groups (either VG21306 or control VG2003 viruses or vehicle). Serum and tumor of both CEACAM6-positive and -negative A549 cell-bearing mice treated with 5 x 10^7^ PFU/mouse of VG21306 candidate via intratumoral injection were harvested at 24 hours, 48 hours, and 14 days post treatment. The concentration of the engager in both the tumor and serum were measured using ELISA. In another experiment, mice were burdened with CEACAM6-negative HepG2 tumor cells and intratumorally injected with 1 x 10^8^ PFU/mouse of VG21306 and the serum and tumors were collected at 48 hours.

## Results

### UCHT-2-03 engager expression and characterization

SPR analysis demonstrated the UCHT-2-03 bispecific antibody showed very strong and specific binding to human CD3 and human CEACAM-6 with affinities (*K*_D_s) of 13.3±0.7 and 407±4.6 pM, respectively (**Figure 1A**). UCHT-2-03 did not bind to an irrelevant protein EGFR.

**Figure 1:**
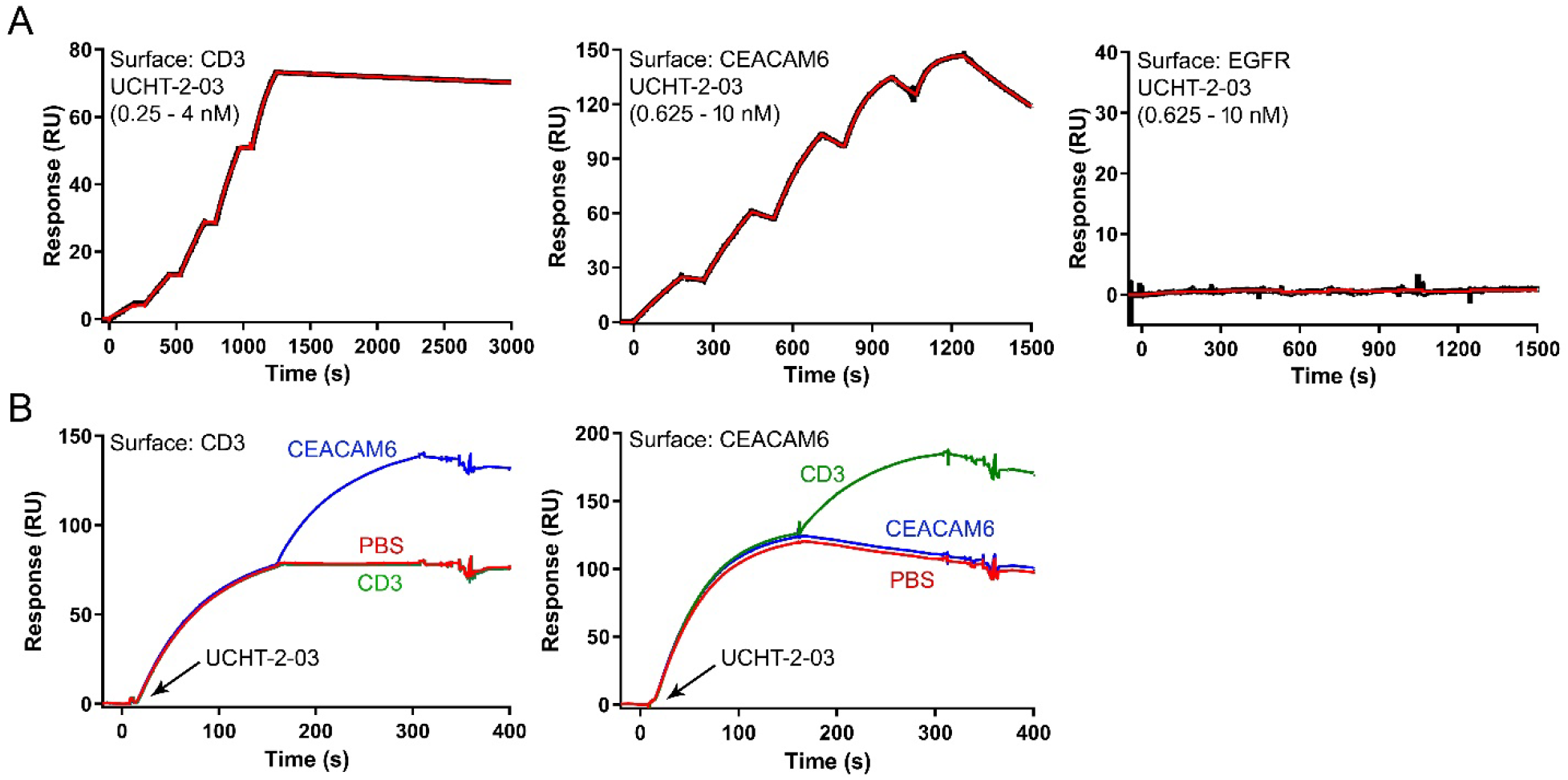
Characterization of T-cell engager binding affinity and co-engagement using SPR. **(A)** UCHT-2-03 binding affinities for immobilized CEACAM6 and CD3δ/ε heterodimer were determined by flowing UCHT-2-03 at concentrations ranging from 0.25 – 4 nM (CD3 surface) and 0.625 – 10 nM (CEACAM6 and EGFR surfaces) and fitting to a 1:1 binding model. **(B)** SPR sensorgrams demonstrating the simultaneous binding of both arms of UCHT-2-03. (*Left*) UCHT-2-03 (10 nM) injected over CD3, followed by injection of CEACAM6 (10 nM, blue line), CD3 (10 nM, green line), or HBS-EP buffer (red line). (*Right*) UCHT-2-03 (10 nM) injected over CEACAM6, followed by injection of CEACAM6 (10 nM, blue line), CD3 (10 nM, green line), or HBS-EP buffer (red line). 1: first injection point; 2: second injection point.

UCHT-2-03 was also capable of binding both antigens simultaneously. UCHT-2-03 was pre-bound to immobilized human CD3 or immobilized humanCEACAM-6 and was shown to bind the second antigen supplied in solution (**Figure 1B**).

### UCHT-2-03 engager cross-reactivity characterization

To evaluate the cross-reactivity of the VG21306 payload (UCHT-2-03 engager) on normal and cancer tissues, recombinant UCHT-2-03 engager was added on multi-organ human cancer or normal tissue microarray slides. Tissues bound UCHT-2-03 then were detected using an HRP-conjugated anti-6X-His-tag antibody. Slides were then stained in the 3, 3’-diaminobenzidine (DAB) chromogenic detection assay. Immunostaining frequency and intensity were given a score between 0 to 4 as described in the Methods section.

Various levels of expression were shown in different tumor tissue sections. The highest level of expression was shown in the pancreas duct adenocarcinoma tissue, stomach adenocarcinoma tissue, and liver compounded hepatocellular carcinoma tissue, all of which had a frequency and intensity score of 4 (**Table 2**). Other tumor tissue sections with both frequency and intensity scores above 3 are listed in Table 1.

**Table 1:** Summary of kinetics analysis of UCHT-2-03 binding to hCD3 and hCEACAM6.

| Surface | Antibody | Rmax | ka (1/Ms) | kd (1/s) | KD (M) | KD (pM) |
| --- | --- | --- | --- | --- | --- | --- |
| h-CD3 | UCHT-2-03 | 83.3 | 1.79E+06 | 2.29E-05 | 1.28E-11 | 13 |
|  |  | 83 | 1.79E+06 | 2.36E-05 | 1.31E-11 | 13 |
|  |  | 83.5 | 1.78E+06 | 2.50E-05 | 1.41E-11 | 14 |
| h-CEACAM-6 | UCHT-2-03 | 171.1 | 2.09E+06 | 8.40E-04 | 4.02E-10 | 402 |
|  |  | 160.2 | 2.08E+06 | 8.54E-04 | 4.11E-10 | 411 |
|  |  | 152.8 | 2.07E+06 | 8.45E-04 | 4.08E-10 | 408 |

\*Theoretical Rmax was 203 RU for CD3 surface and 220 RU for CEACAM6 surface

**Table 2:** Tumor Tissue Immunostaining with Frequency and Intensity Scores Above 3

| Tissue | Microarray Panel | Frequency | Intensity |
| --- | --- | --- | --- |
| A6 Pancreas duct adenocarcinoma | BC001157a | 4 | 3 |
| B6 Pancreas duct adenocarcinoma | BC001157a | 4 | 3 |
| C6 Pancreas duct adenocarcinoma | BC001157a | 4 | 4 |
| E8 Stomach adenocarcinoma | BC001157a | 4 | 4 |
| E9 Lung bronchioloalveolar carcinoma (muoinous) | BC001157a | 4 | 3 |
| F3 Colon adenocarcinoma | BC001157a | 3 | 4 |
| F6 Rectum adenocarcinoma (sparse) | BC001157a | 3 | 4 |
| F9 Lung adenosquamous carcinoma | BC001157a | 4 | 3 |
| G4 Liver compounded hepatocellular carcinoma | BC001157a | 4 | 4 |
| G8 Stomach adenocarcinoma | BC001157a | 3 | 3 |

Although most normal tissues did not show any binding, there were some exceptions where UCHT-2-03 binding was observed in some normal tissues (**Table 3**). Normal spleen tissue sections had a frequency score of 3 and an intensity score of 2. A normal kidney tissue section had a lower level of expression with a frequency score of either 1 or 2 and an intensity score of 1. There was also weak staining in one normal lung tissue section, which had frequency and intensity scores of 1 (**Table 3**).

**Table 3:** Normal Tissues Showing Some Level of Binding

| **Tissue** | **Microarray Panel** | **Frequency Score** | **Intensity Score** |
| --- | --- | --- | --- |
| C5 Lung tissue | BN961a | 1 | 1 |
| D3 Spleen tissue | BN961a | 3 | 2 |
| D4 Spleen tissue | BN961a | 3 | 2 |
| E3 Kidney tissue | BN961a | 2 | 1 |
| E4 Kidney tissue | BN961a | 1 | 1 |

### In vitro recruitment of T cells by UCHT-2-03 T-cell engager

The results show that T cells can be engaged by VG21306-expressed CD3/CEACAM6 T-cell engager in the presence of CEACAM6-expressing A549 cells, but not in CEACAM-negative A549 cells. To visualize the binding of UCHT-2-03 engager with CEACAM6-positive human lung cancer cells (A549) and purified human T cells, we stained A549 or CEACAM6 knockout (KO) A549 cells and T cells with red dye (CellTracker™ Red) and green dye (CellTracker™ Green), respectively. Tumor and T cell co-cultures were then incubated at 37°C with either DPBS (vehicle) or backbone virus VG2003 (without UCHT-2-03 engager) and VG21306. A549 cells with CEACAM6 KO A549 cells were used as the comparison group. An abundance of T cells surrounding CEACAM6-positive A549 cells was observed, while very few T cells were evident surrounding CEACAM6-negative A549 cells. HSV-1 staining confirms the infection of A549 cells (**Figure 2**).

**Figure 2:**
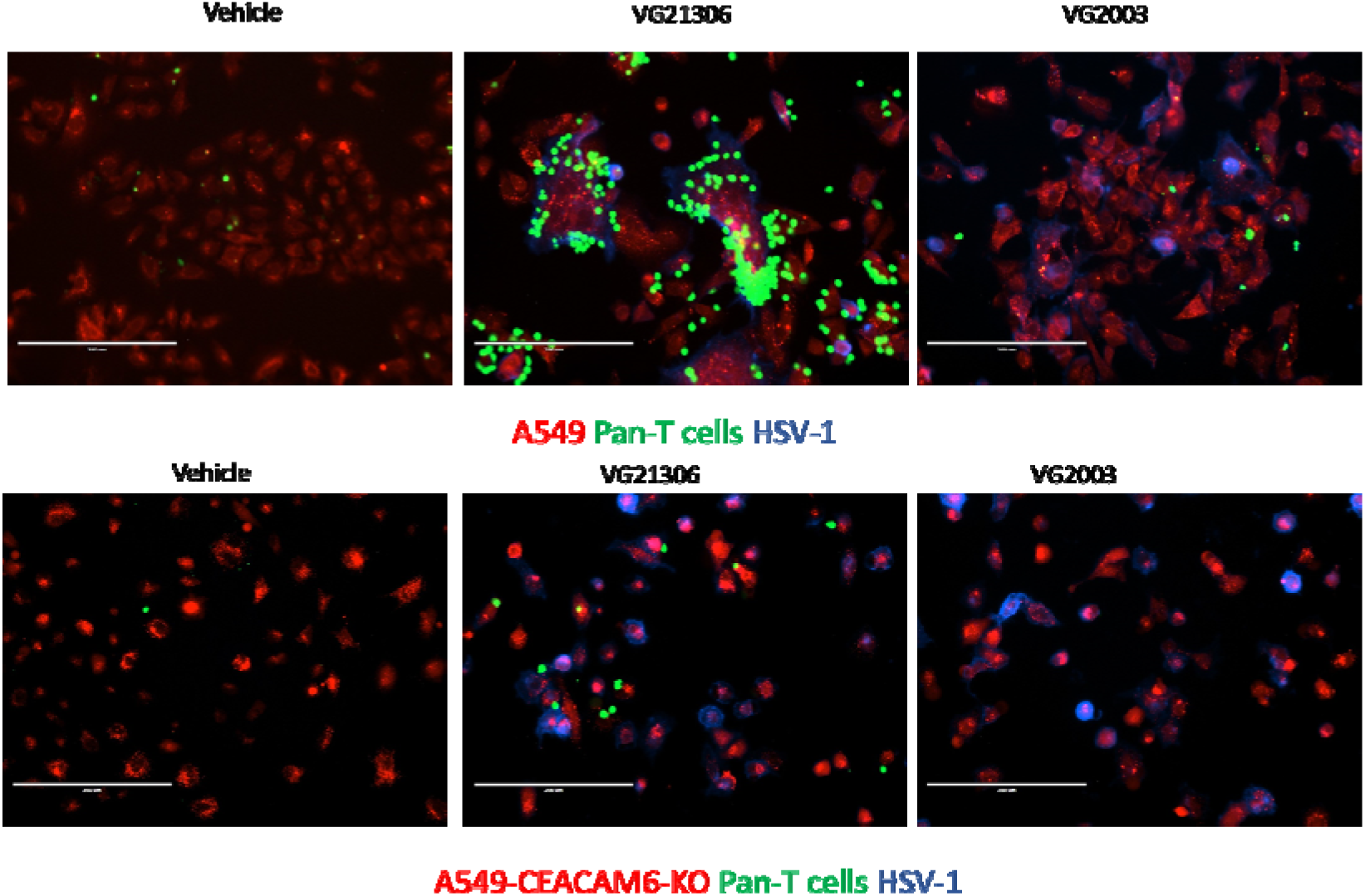
Effect of VG21306 in A549-CEACAM6-KO and T cell co-culture system in vitro. Indicated cells were stained with CellTracker™ Red, seeded into a 24-well plate, then incubated at 37°C overnight. Pan-T cells were then stained with CellTracker™ Green and co-cultured with previously seeded A549-CEACAM6 KO cells. Cells were then incubated with either DPBS or VG21306 or VG2003 virus at an MOI of 2 at 37°C for 18 - 20 hours. Cells were then washed twice with DPBS and fixed with 4% PFA. Fixed cells were then stained with anti-HSV-1 antibody to detect virus-infected cells. Imaging (20X) was performed using a fluorescence microscope.

### In vitro killing effect of UCHT-2-03 T-cell engager

To determine if the UCHT-2-03 T-cell engager effectively kills different CEACAM6-positive tumor cells, an in vitro assay was carried out whereby human T cells from healthy donors were co-incubated with CellTrace™ Far Red dye-labelled A549 lung cancer and MDA-MB-468 breast cancer cells in the presence or absence of engager. After incubating at 37°C for 48 hours, cells were harvested and incubated with the viability dye 7AAD (**Figure 3**). Approximately 50% and 70% more A549 and MDA-MB-468 cells were killed in the presence of engager compared to the background level of cell killing in the absence of engager (13% in A549 cells and 24% in MDA-MB-468 cells). These studies demonstrate that a variety of tumor cell types with CEACAM6 expression can be killed by engager-activated T cells.

**Figure 3:**
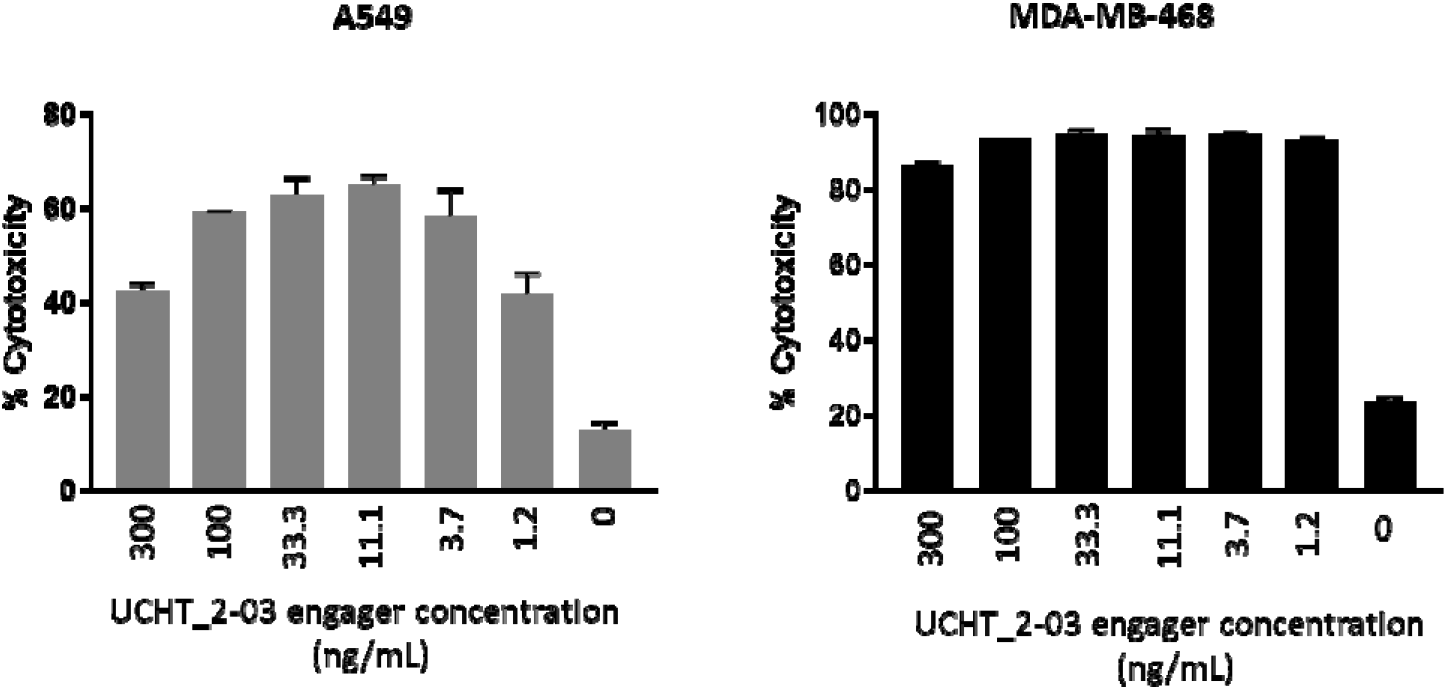
UCHT-2-03 T-cell engager activates T cell killing against CEACAM6^+^ tumor cells. A549 and MDA-MB-468 tumor cells were labelled with CellTrace™ Far Red and incubated with healthy human T cells and different doses of engager, incubated at 37°C then analyzed 48 hours later for the percentage of dead tumor cells. Values for tumor cell cytotoxicity represents the average ± SD of two replicates.

### T-cell engager-armed oncolytic HSV enhances tumor cytotoxicity

To evaluate whether UCHT-2-03 engager expressed as virus payload enhances tumor-cell killing, we co-cultured healthy PBMCs with A549 tumor cells infected with VG21306 virus (expressing human IL-12 and IL-15/IL-15Ra, and T-cell engager as payloads), VG2003 virus (expressing human IL-12 and IL-15/IL-15Ra payloads), or VG1905 backbone virus (no payload expression) at MOI=0.1 at 37°C for 5 days. We also performed a co-culture experiment in the presence of 10 ng/mL of purified CD3/CEACAM6 engager as a control. A549 cell killing was monitored and analyzed in real time by impedance. As shown in **Figure 4**, both control VG1905 and VG2003 viruses induced 20% higher tumor-cell killing compared to the co-culture without virus; however, a significant increase (60% of increased cytolysis) in tumor cytotoxicity was observed when VG21306 virus was used, suggesting an enhanced cytotoxic role of CD3/CEACAM6 T-cell engager against tumor cells.

**Figure 4:**
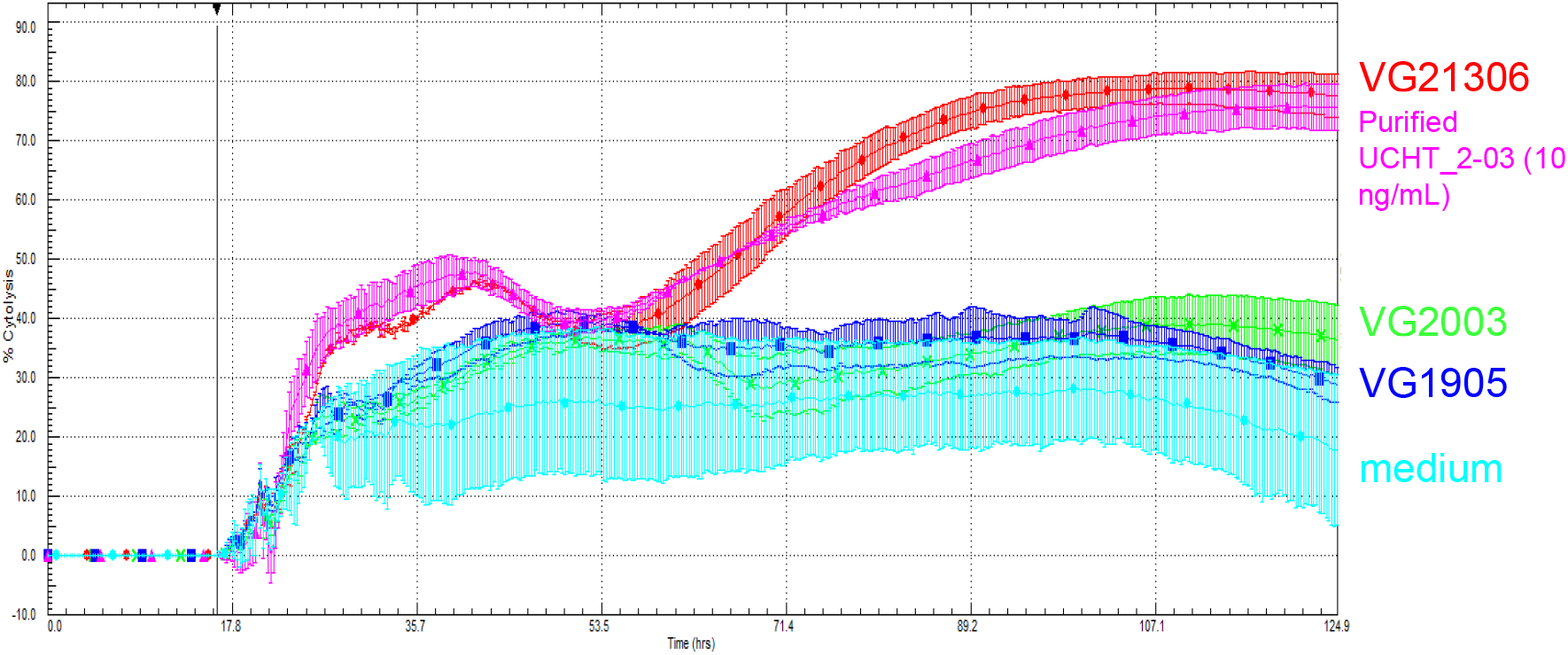
Engager-mediated oncolysis enhances tumor cell cytotoxicity. A real-time xCELLigence-based cytotoxicity assay was used to evaluate the lysis of A549 cells over a 120-hour period. Tumor cells were seeded and infected with different viruses at t0 and incubated at 37°C. Healthy human PBMCs were added 18 hours later. Impedance at well bottoms was measured every 15 minutes up to 120 hours and normalized to baseline impedance values with A549 cells co-cultured with medium only.

### In vivo T cell activation by UCHT-2-03 engager

To examine the in vivo effect of UCHT-2-03 T-cell engager on T-cell activation, a humanized NSG mouse model was used. NSG mice were subcutaneously transplanted with A549 tumor cells and human PBMCs and intratumorally injected with engager-expressing VG21306 virus, control VG2003 virus, or vehicle. As displayed in **Figure 5**, although we did not observe increased T-cell expansion in mice treated with VG21306 compared with the other groups, VG21306 virus injection showed an increased CD69^+^ T cell percentage in tumors. This result suggests CD3/CEACAM6 T-cell engager activates T cells.

**Figure 5:**
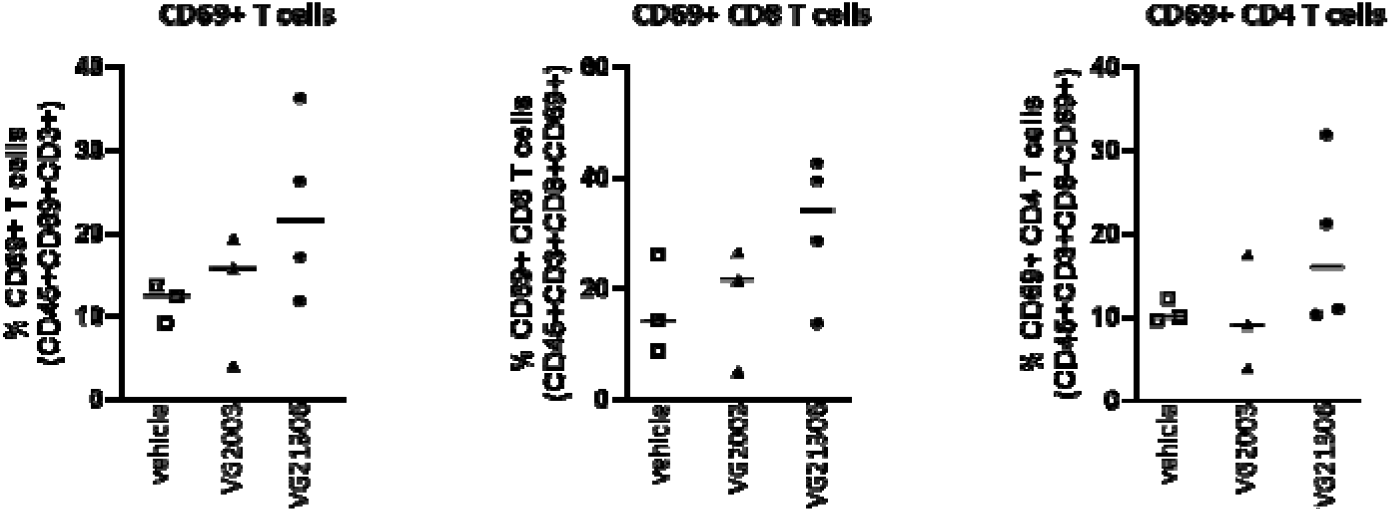
VG21306-enhanced T-cell activation in vivo. A549 cells were mixed with human PBMCs from a healthy donor and transplanted into the right flanks of NSG mice. A total of 5 x 10^7^ PFU of VG21306 or VG2003 virus was injected intratumorally once the tumor size reached 100 mm^3^. Tumor samples were isolated at 3 days after treatment and percentages of activated T cells were identified by staining with human CD45, CD3, CD4, CD8, and CD69 antibodies and analyzed by flow cytometry.

### Leaking of engager payload in different tumor models

Since one of the drawbacks of using T-cell engagers for systemic delivery was on-target, off-tumor toxicity, and to ensure that the payload engager expressed by the VG21306 virus was restricted to the injected tumors (not leaked to the blood), we performed an in vivo test where the virus was injected intratumorally into different types of tumors, including CEACAM6-positive and -negative types. As shown in **Figure 6**, engager expression was detected in the tumor tissue up to 14 days after the virus injection; however, no detectable amounts were measured in the serum. These results were not related to CEACAM6 expression by the tumor.

**Figure 6:**
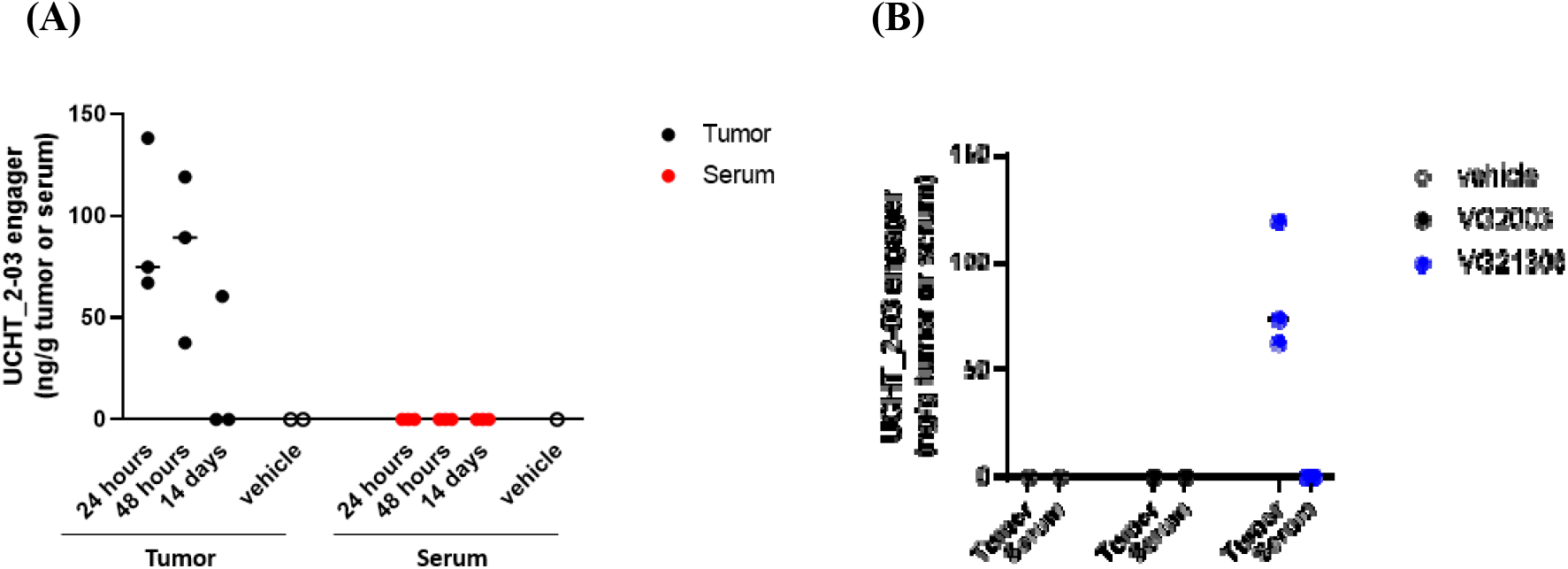
Detection of UCHT-2-03 engager in tumor and circulating blood. CEACAM6-positive A549 tumor (A) or CEACAM6-negative HepG2 tumor (B) were implanted in the flank of athymic nude mice and subsequently received intratumoral injection of vehicle, VG2003 virus, or VG21306 virus. At indicated time points in the figure, tumor samples were harvested and homogenized to obtain protein lysates. In addition, mouse serum was isolated from whole blood samples. The concentration of UCHT-2-03 engager in tumor and serum samples was assessed by ELISA.

### Bystander killing of CEACAM6-negative cells

To determine the effect of the UCHT-2-03 engager on cells that do not express the CEACAM6 antigen, we’ve designed an experiment where both CEACAM6-positive or -negative cells were cultured separately or in co-culture, then the engager and pan-T cells were applied to the culture. After incubation at 37°C for 48 hours, the cell killing as well as granzyme B and IFN-γ secretion by the T cells were measured. The results indicate that when the tumor cells were grown separately, and in the presence of both the engager and T cells, only the tumor cells expressing CEACAM6 (BxPC3) were killed (**Figure 7a**), while CEACAM6-negative cells (HepG2) were not affected by the treatment (**Figure 7b**). However, when the two different types of tumor cells were co-cultured, the CEACAM6-negative tumor cells showed significant killing. Moreover, the secretion of both granzyme B and IFN-γ was induced only when the CEACAMpositive cells were present, either alone or in co-culture with CEACAM6-positive cells. No significant amounts were detected with CEACAM6-negative cells alone (**Figure 7c and 7d**).

**Figure 7:**
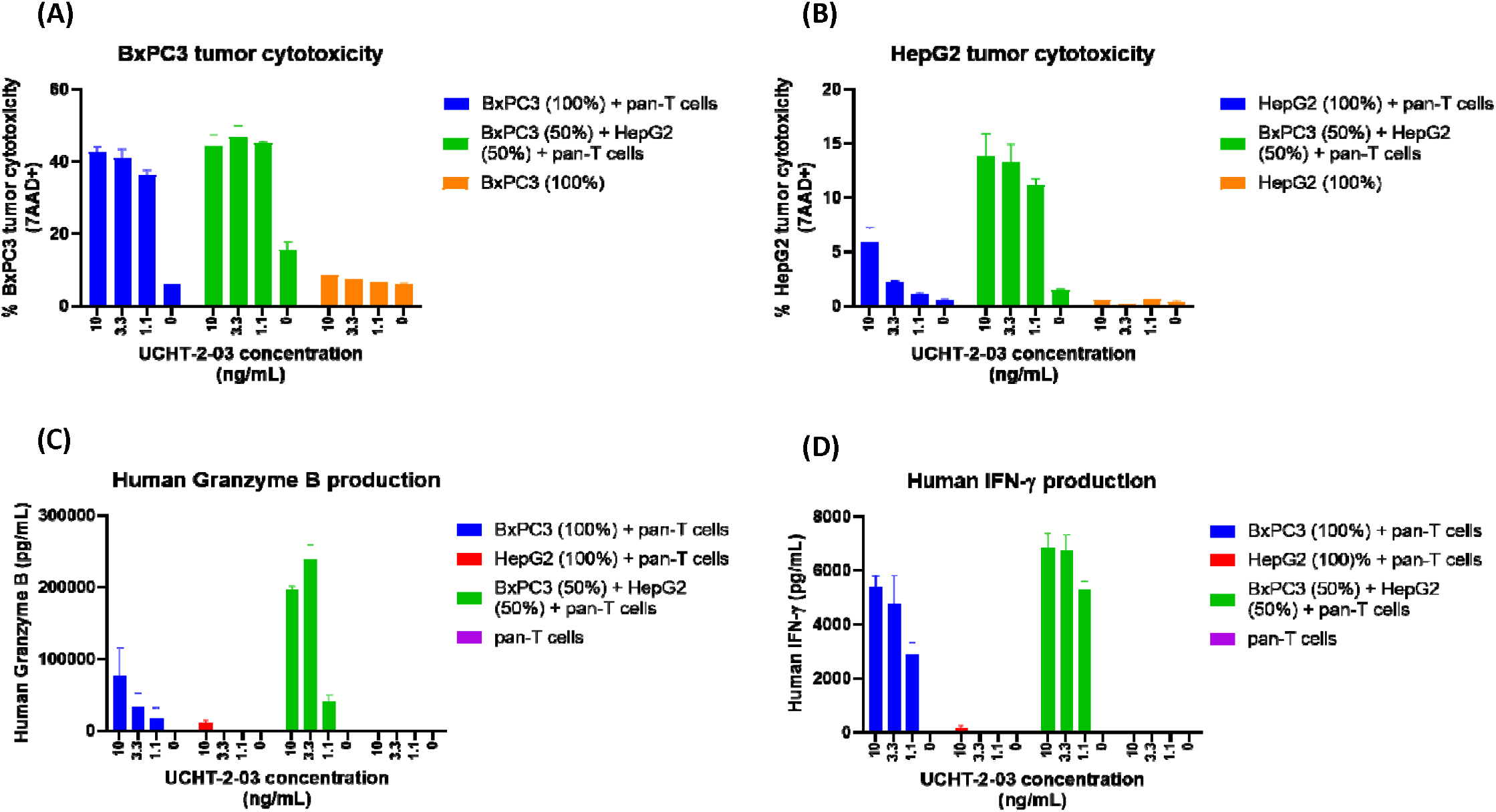
Detection of bystander killing using UCHT-2-03 engager. The killing efficacy of CEACAM6-positive BxPC3 tumor cells and CEACAM6-negative HepG2 tumor cells was tested using viability dye 7AAD (A and B, respectively). The secretion of granzyme B and IFN-γ was measured using ELISA (C and D).

### UCHT-2-03 killing using exhausted T cells

To test whether the immunosuppressive conditions in the tumor microenvironment will impact the killing of tumor cells via exhausted T cells, we performed a test in which pan-T cells were exposed to continuous stimulation for 9 days, resulting in exhausted phenotype (high expression of PD-1, TIM3, TIGIT and LAG3) (**Figure 8a**). The exhausted T cells were used in a killing assay using the UCHT-2-03 engager. The tumor-cell killing was compared to that induced by freshly thawed (non-exhausted) T cells from the same donor. The results indicate that regardless of the condition of the T cells they were still able to induce cell death on tumor cells in the presence of UCHT-2-03 engager (**Figure 8b**).

**Figure 8:**
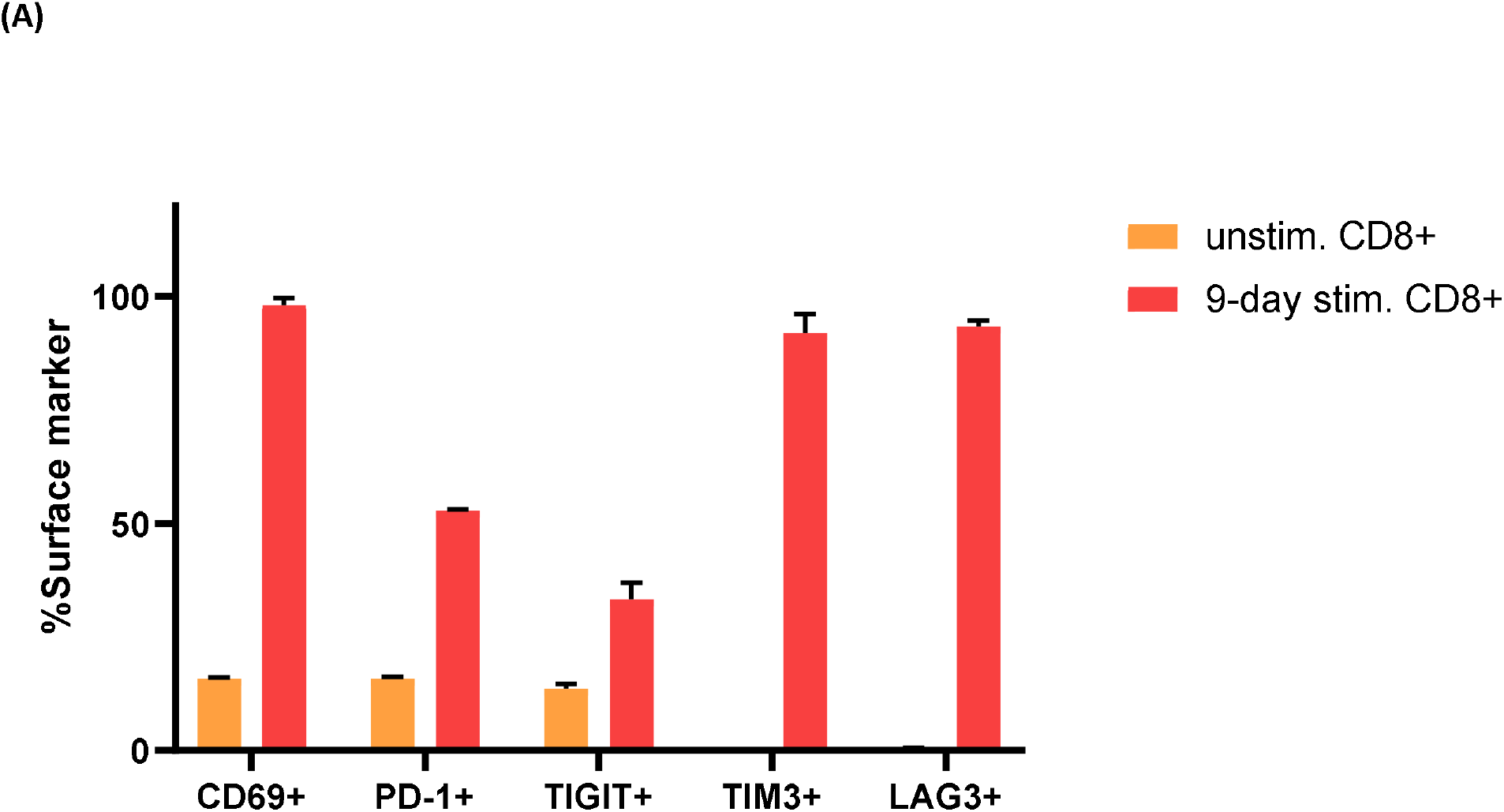
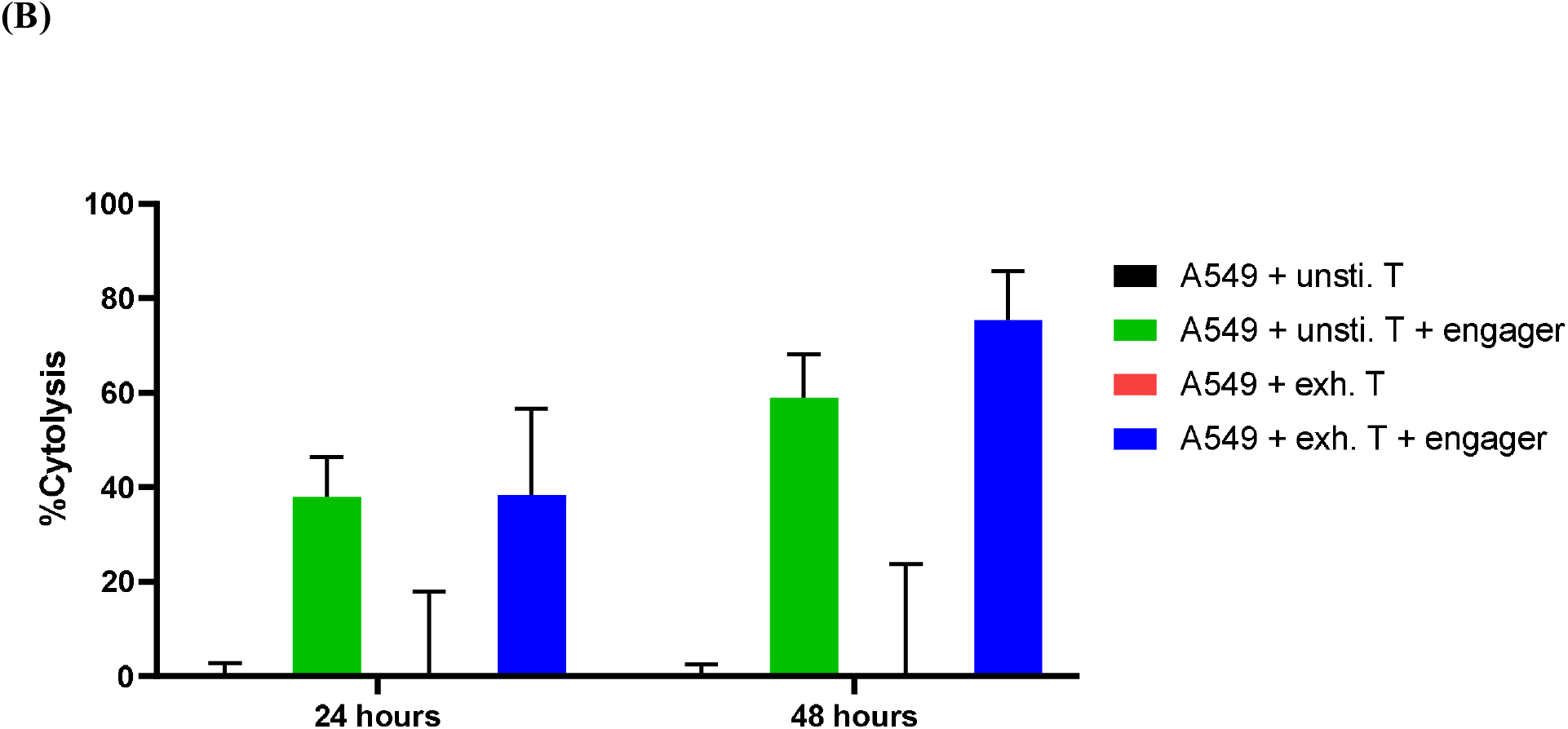
UCHT-2-03 engager induced killing using exhausted T-cell. Pan T-cells were exposed to continuous stimulation, resulting in higher expression of stimulation and exhaustion markers (A). The exhausted T-cell, along with unstimulated T-cells, were used in a killing assay using BxPC3 tumor cells and UCHT-2-03 engager (B).

## Discussion

Oncolytic viruses (OVs) are a powerful tool for fighting cancer. Besides their ability to directly attack and lyse cancer cells, they have also demonstrated an impressive ability to induce a potent and lasting memory immune response against tumors [30][28]. Oncolytic virotherapy relies on tumor cell-specific changes associated with the hallmarks of cancer, including increased receptor expression, impaired antiviral response, and altered metabolism [32]. OVs can be engineered to express immunotherapeutic transgenes directly at the site of infection, achieving high local concentrations while preventing systemic side effects [33]. Furthermore, oncolysis can render immune-excluded and immunosuppressed tumors sensitive to otherwise unsuccessful strategies such as immune checkpoint inhibition. For this reason, it has been argued that the future of immunotherapy will include OVs that function as multiplexed immune-modulating platforms expressing factors such as immune checkpoint inhibitors, tumor antigens, cytokines, and T-cell engagers [33].

Previously, we developed a novel HSV-1-based OV that is based on the TTDR platform [30]. In the TTDR platform, we enhanced the potency of the virus by regulating viral genes ICP27 and ICP34.5, and thus avoided attenuating the virus to achieve a safe OV [12]. In this work, we have introduced a novel payload to further enhance the anticancer activity of VG2025 [30]—a bispecific T-cell engager (BiTE) targeting T cells at tumor cells expressing CEACAM6 antigen.

T-cell engagers have been utilized effectively to treat and eliminate liquid cancers (i.e, leukemias) by targeting specific surface markers. The use of the same concept to target solid tumors has been hindered, mostly by two factors: toxicity and inability to reach the tumor site efficiently [34][35]. In this work, we tried to tackle both issues by delivering and limiting the expression of the T-cell engager to the tumor site to maximize the efficacy and reduce the possibility of inducing a systemic toxic immune response. Moreover, locally expressed payload will circumvent the problem of reaching the tumor site when delivered systemically. We have demonstrated the expression of a functional engager that was able to both engage and activate T cells in vitro and in vivo (**Figures 3 and 5**). The results from in vivo models demonstrated that very little—if any—of the expressed engager was detected in the blood, while nanogram levels were detected in the injected tumors up to 14 days post virus injection (**Figure 6**). This work is critical in demonstrating the safety of this approach and showing that none of the expressed engager would leak to the blood.

As most tumors in real life are heterogeneous, it is not expected that 100% of the tumor mass will express a certain tumor antigen. To study the effect of the expressed engager on cells not expressing the tumor antigen, we’ve demonstrated that even when tumor cells don’t express the tumor antigen (CEACAM6), the presence of even a fraction of the cells that express the antigen was enough to induce the killing of non-engaged cells (**Figure 7**). Moreover, the engager was also effective in inducing cell death even when the T cells were exhausted (**Figure 8**).

In conclusion, VG21306 is a novel OV designed to target CEACAM6-expressing tumors via a T-cell engager payload. Besides offering the full therapeutic benefits of an OV, this new OV offers extra efficacy by expressing the T-cell engager in the tumor microenvironment.

## Author Contributions

Conceptualization, **Y.M**., W.J., and W.L..; methodology, **Y.M., J.D**., I-F.L., **Z.D**., W.L. O.T.; G.L., D.C., G.H., and H.F. formal analysis, Y.M., Z.D., I-F.L., W.L., G.H., and H.F.; writing—original draft preparation, Y.M.; writing—review and editing, Y.M., I-F.L., Z.D., G.H., W.J. All authors have read and agreed to the published version of the manuscript.

## Funding

This research received no external funding

## Conflicts of Interest

**Y.M**., I-F.L., X.L., Z.D., J.D., D.C., G.L., O.T., and W.J. are employees of Virogin Biotech, a company focusing on developing oncolytic virotherapy.

## Acknowledgments

We thank Marianne Chomiak for critical reading of the manuscript. We also thank Ceren Gulhan, and Sehyun Kim for technical assistance.

## Notes

### Summary of Updates

I wanted to update the affiliation of the last author to match the PDF file, but it was already fixed, so no changes were made. Sorry for the inconvenience.

